# Evaluation of lung VEGF-A transduction during hyperoxia-induced injury in rats

**DOI:** 10.1101/306381

**Authors:** William Raoul, Vanessa Louzier, Saadia Eddahibi, Serge Adnot, Bernard Maitre, Christophe Delclaux

## Abstract

**Background:** Since Vascular Endothelial Growth Factor (VEGF) is a main factor for endothelial survival, we evaluated whether VEGF transduction could ameliorate hyperoxia induced injury, which is associated with predominant endothelial injury.

**Methods and Results:** Transduction (induced 48 hours before hyperoxic exposure) using adenoviral vector (Ad.) for VEGF (10^10^ viral particles [VP]) increased moderately survival under hyperoxia (fraction of inspired oxygen [FIO_2_] >95%) as compared with Ad.Null (10^10^ VP) transduction, whereas VEGF transduction with a lower dose (5.10^9^ VP) had no effect. After 48 hours of hyperoxia, Ad.VEGF transduction increased lung VEGF concentration, prevented the diffuse loss of capillary bed and induced patchy areas of endothelial cell proliferation (CD31 immunostaining) with interstitial inflammatory cell recruitment as compared to Ad. Null transduction. Hyperoxia was associated with diffuse apoptosis that was inhibited only in patchy areas of endothelial proliferation under VEGF transduction. Hyperoxia-induced alveolar inflammation was similar with Ad.Null and Ad.VEGF. Under normoxia, the high dose of VEGF transduction induced diffuse alveolar inflammation whereas the low dose did not suggesting a pro-inflammatory effect of VEGF that may have participated to increased survival under hyperoxia.

**Conclusions:** We demonstrate that lung VEGF-A transduction despite inhibition of the loss of capillary bed has a marginal effect of on animal survival during hyperoxia-induced injury.

## Background

Increased permeability and interstitial and pulmonary edema are prominent features of acute lung injury/acute respiratory distress syndrome (ARDS), and Vascular Endothelial Growth Factors (VEGFs) and their receptors have been implicated in the regulation of vascular permeability in many organ systems, including the lung [1]. On the other hand, a decrease in lung capillary density has clearly been demonstrated during ARDS [2], which may lead to an increase in alveolar dead space, a factor associated with increased lethality [3]. VEGF has been demonstrated to be a major physiological survival factor for endothelium [4], which is abundantly secreted in healthy lung [5]. We and others evidenced low levels of VEGF in the lungs of patients with ARDS [6, 7]. This VEGF decrease may have contributed to vascular injury inasmuch as we subsequently demonstrated a negative relationship between VEGF concentration in lung homogenate and the number of endothelial cells undergoing apoptosis in lung tissue of patients suffering from ARDS [8]. VEGF (or VEGF-A) is a highly conserved, dimeric, heparin-binding glycoprotein (molecular weight 46 kDa). At least four different VEGF transcripts resulting from alternate splicing of a single gene have been identified in human cells. VEGF_121_ and VEGF_165_ are secreted in soluble form, whereas VEGF_189_ and VEGF_206_ remain cell-surface associated or are primarily deposited in the extracellular matrix. VEGF seems to specifically affect endothelial cell growth, survival, and permeability. In the lung, VEGF is expressed primarily by epithelial cells and macrophages. The biological activity of VEGF is dependent on interaction with specific receptors (VEGF-R1, 2, 3), which are expressed not only by endothelial cells but also by activated macrophages and alveolar type II epithelial cells. Endothelial survival is mediated via VEGF-R2/KDR [9]. The aim of this study was to determine whether VEGF is a main factor determining endothelial and animal survival during acute lung injury. To this end, we used VEGF adenoviral-mediated transduction during hyperoxia-induced injury inasmuch as hyperoxia has been associated with predominant endothelial injury [10].

## Materials and Methods

### Design of the study

#### Preliminary set of experiments

a time course study of lung VEGF expression after adenoviral-transduction has been previously published by our laboratory [11]. Based on these results, adenoviral transduction was conducted 48 hours before hyperoxia exposure inasmuch as significant levels of human VEGF are already expressed in lungs of rats in accordance with the results obtained by Kaner and colleagues [12].

#### First set of experiments

the survival of rats that received either Ad.VEGF or Ad.Null was determined under hyperoxia (fraction of inspired oxygen [FIO_2_] >95%). Two doses (expressed as viral particles [VP], 5.10^9^, 10^10^) of VEGF-transduction were evaluated.

#### Second set of experiments

rats challenged with either Ad.VEGF or Ad.Null were sacrificed on the second day of exposure under hyperoxia (FIO_2_ >95%) in order to assess lung histopathology, edema, number of alveolar inflammatory cells and lung VEGF concentration. This time point was chosen based on the time-course study under hyperoxia of lung histopathology published by Crapo and colleagues demonstrating significant endothelial injury whereas no mortality is still observed [10]. In this set of experiments, additional rats challenged with either Ad.VEGF or Ad.Null were evaluated under normoxia at similar time point in order to evaluate the respective effects of Ad. VEGF and hyperoxia on indexes of lung injury.

### Recombinant Adenovirus Vectors

All Ad vectors were Ela^-^, partial Elb^-^, partial E3^-^ based on the Ad5 backbone, containing a transgene (AdVEGF165) or no transgene (AdNull) under control of the cytomegalovirus immediate-early promoter/enhancer in the E1a position [12]. AdVEGF165 expresses the human VEGF165 cDNA. All adenovirus vectors were propagated in 293 cells, purified by CsCl gradient centrifugation, dialyzed, and stored at 80°C. Adenovirus vectors were provided by the Vector Core of the University Hospital of Nantes supported by the Association Française contre les Myopathies (AFM).

### Animals and Delivery of Adenovirus Vectors to the Lungs

Male Wistar rats (200 to 220 g body weight) were used for all studies. All animal care and procedures were in accordance with institutional guidelines. Ad.VEGF, or Ad.Null as the control, was diluted before use with sterile saline (Phosphate buffer saline, PBS purchased from Gibco), pH 7.4, in a final volume of 120 μl. Rats were anesthetized with intraperitoneal ketamine (7 mg/100 g) and xylazine (1 mg/100 g). Intratracheal instillation of 120 μl/rat of diluted Ad.VEGF or Ad.Null was performed using a standard procedure, as previously described [11].

### Animal Exposure to Hyperoxia

Two days after intratracheal Ad.VEGF (5.10^9^ and 10^10^ VP) or Ad.V152 (10^10^ VP) administration, the rats were exposed to >95% oxygen in a Plexiglas hyperoxia chamber. The flow of gas was kept constant at 8 L/min. The inspired fraction of oxygen was checked using a gas analyzer (Radiometer ABL330, Copenhagen) at the end of each experiment. The rats had free access to food and water and were housed at room temperature.

### Bronchoalveolar lavage and differential cell count

After intraperitoneal administration of pentobarbital (60 mg/ kg), the abdominal aorta was cut and a median sternotomy was performed. The left lung was used for bronchoalveolar lavage (BAL), which was carried out by flushing the lung with sterile, pyrogen-free, physiological saline via a tracheal cannula (one 5 ml and two 2.5 ml aliquots). The recovered fluids were pooled. The total number of cells was counted using a standard hemocytometer. Cytospin preparations were made using a Shandon 3 cytocentrifuge (Shandon, France). The cells were fixed and stained with Diff-Quick kit (Dade Behring, France). Differential counts on 200 cells were done using standard morphological criteria.

### Tissue Harvesting and Processing

After BAL procedure, the lung tissue was inflated with paraformaldehyde 4% and fixed overnight in paraformaldehyde. Tissues were then dehydrated through serial ethanol concentrations of 70%, 95% and 100%. Tissues were then processed through xylene and embedded in paraffin. Five micron-sections were cut and placed on positively charged slides (Super Frost Plus). To perform CD31 immunohistochemistry, another series of lungs was fixed in Tissue-Tek (Sakura) /PBS (1:1) and frozen in liquid nitrogen. Five to six-micron-thick cryosections were then performed with cryostat (JUNG CM300, Leica).

### Quantification of Lung Wet/Dry Weight Ratio

The right lung was used for edema assessment and tissue homogenate preparation. A lung sample was immediately weighed and then placed in a dessicating oven at 65°C for 48 h, at which point dry weight was achieved. The ratio of wet/dry weight was used to quantify lung water content, an index of lung edema. The remaining tissue was fixed in liquid nitrogen to allow subsequent homogenate preparation.

### VEGF concentration in lung homogenates

A lung tissue specimen was thawed and homogenized in lysis buffer (10 mg/ml) containing 0. 5% Triton X100, 150 mM NaCl, 15 mM Tris, 1 mM CaCl, and 1 mM MgCl (pH 7.4). Homogenates were incubated on ice for 30 min, then centrifuged at 12 000 g for 20 min at 4°C. Supernatants were collected then stored at −80°C for assays. The concentration VEGF was determined using an human sandwich enzyme-linked immunosorbent assay (ELISA) according to the manufacturer’s instructions (R&D Systems, France). We and others previously demonstrated that this human VEGF ELISA kit cross reacts with rat VEGF [5, 13].

### Immunohistochemistry: CD31 staining

Purified monoclonal mouse anti-rat CD31 (BD Pharmingen) antibody was used to quantify the microvascular bed in our experiments. Briefly, frozen slides were allowed to come to room temperature and were then fixed with paraformaldehyde 1% in PBS, rinsed and put in ethanol / acetic acid (2:1) for 5 min at −20°C. Slides were incubated in 3% H_2_O_2_ for 15 min to quench endogenous peroxidase and rinsed. Non specific sites were blocked with Goat serum 2% in PBS tween 0.05% at RT. Monoclonal mouse anti-rat CD31 antibody was incubated (diluted 1/100 in PBS/BSA 1%) for 2 hours at RT in humidified chamber. After rinsing, second antibody goat anti-mouse IgG from Sigma (diluted 1/100) was processed for 30 min in humidified chamber at RT. Slides were labeled with ExtrAvidin peroxidase for 30 min. 3,3’- diaminobenzidine tetrahydrochloride dihydrate (DAB, Sigma) was used to visualize labeled antibody.

### TUNEL staining

In situ detection of apoptotic cells was evaluated by terminal deoxynucleotidyl transferase (TdT)-mediated dUTP nick end labeling (TUNEL) using a commercially kit (ApopDETECT, QBiogene, France), according to the manufacturer’s protocol. Briefly, after paraffin removal in xylene, the sections were rehydrated and exposed to proteinase K (Promega, USA). Endogenous peroxidase was quenched with 3% H_2_O_2_. Residues of digoxigenin-nucleotide were catalytically added to the DNA by TdT to the 3’-OH ends of DNA at 37°C for 1 hour. The reaction was visualized by peroxidase-conjugated anti-digoxigenin antibody and 3,3’- diaminobenzidine, DAB, which colored nuclei into dark brown. The sections were counterstained by 0.5% methyl green. Percentage of TUNEL-positive cells was evaluated over 1000 cells in multiple fields in 3 sections per condition (control FIO_2_ 21 %, Ad.Null 10^10^ VP and AdVEGF 10^10^ VP in normoxia). Only cells that showed typically morphologic apoptosis features (blebbing cytoplasmic membrane, condensed chromatin and nucleus shrinkage) were taken into account.

### Histological analyses and quantification

All the histological evaluations were performed in blind fashion. Inasmuch as initial histological evaluation showed patchy areas with cellular proliferation, all subsequent quantitative analyses were performed with and without selection of areas of obvious cellular proliferation. This was justified by the fact that quantitative analysis of these areas could induce a bias, our aim was to evaluate whether adenoviral transduction had a biological effect in the whole lung (especially in areas without cellular proliferation). The inflammatory response was assessed using a previously described empiric semi-quantitative scale [14] based on inflammatory-cell type and location (alveoli, bronchi, or blood vessels). The extent of inflammation was scored on a 0-4 scale as follows: 0, none; 1, small scattered areas; 2, <10%; 3, 10-50%; and 4, >50% of section area.

### Statistical analysis

The data are expressed as means ± standard error of the mean (SEM). The significance of differences in continuous variables between control and individual experimental groups was determined using the Mann-Whitney U test or Kruskall-Wallis test as appropriate. Survival curves (Kaplan-Meier plots) were compared using Breslow-Gehan-Wilcoxon test. Statistical significance was defined as P<0.05. Statistical analysis was performed using STATVIEW 5.0 software (Abacus Concepts, Inc, Berkeley, CA).

## Results

### Effect of Ad.VEGF transduction on rat survival under hyperoxia

All the Ad.Null (10^10^ VP) control rats died between 67 h and 92 h of initiation of 95% O2 exposure. Transduction with the low dose of Ad.VEGF (5.10^9^ VP) did not significantly increase survival under hyperoxia whereas transduction with the high dose of Ad.VEGF (10^10^ VP) induced a moderate but significant increase in survival (p<0.05) (figure 1). We mainly focused the remainder of the study on the high dose of Ad.VEGF.

**Figure 1.**
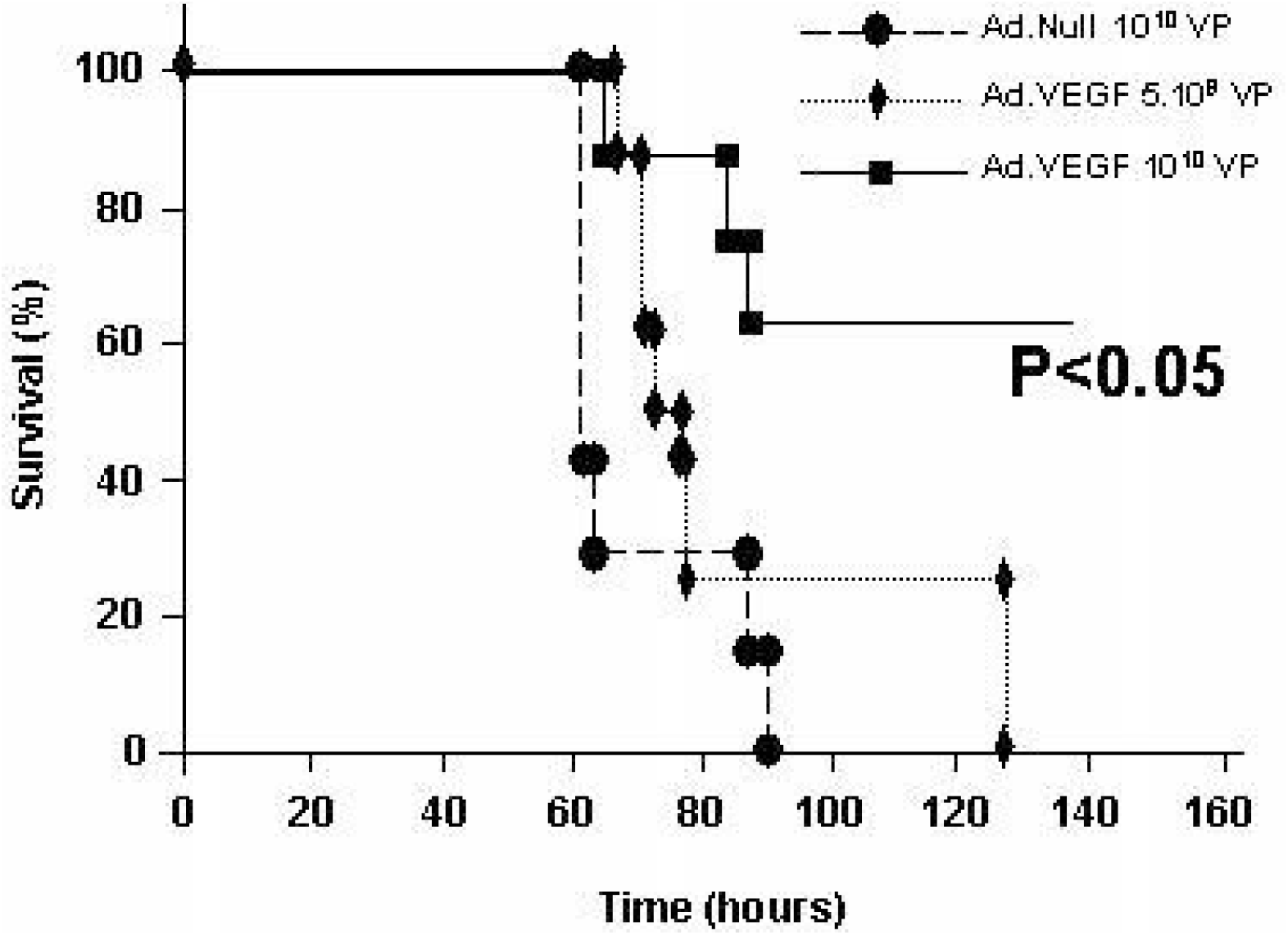
Survival of rats under hyperoxia. Rats were pretreated with Ad.VEGF 5.10^9^ VP (triangles) or Ad.VEGF 10^10^ VP (squares) or with control Ad.Null 10^10^ VP (circles) two days before initiating oxygen exposure. Survival was then assessed (n = 8 for each group).

### Evaluation of VEGF concentration and lung injury indexes after 48 hours of hyperoxia

#### Efficiency of transduction in hyperoxia

##### Quantification of VEGF levels in lung homogenates

In lung homogenates obtained 4 days after Ad.VEGF administration (FIO_2_ 95% during 2 days), a dramatic increase in VEGF concentration was evidenced as compared to Ad.Null transduction. However no dose effect relationship was evidenced (figure 2). There was no statistical difference in VEGF concentration between control lung homogenates under normoxia and Ad.Null homogenates under hyperoxia.

**Figure 2.**
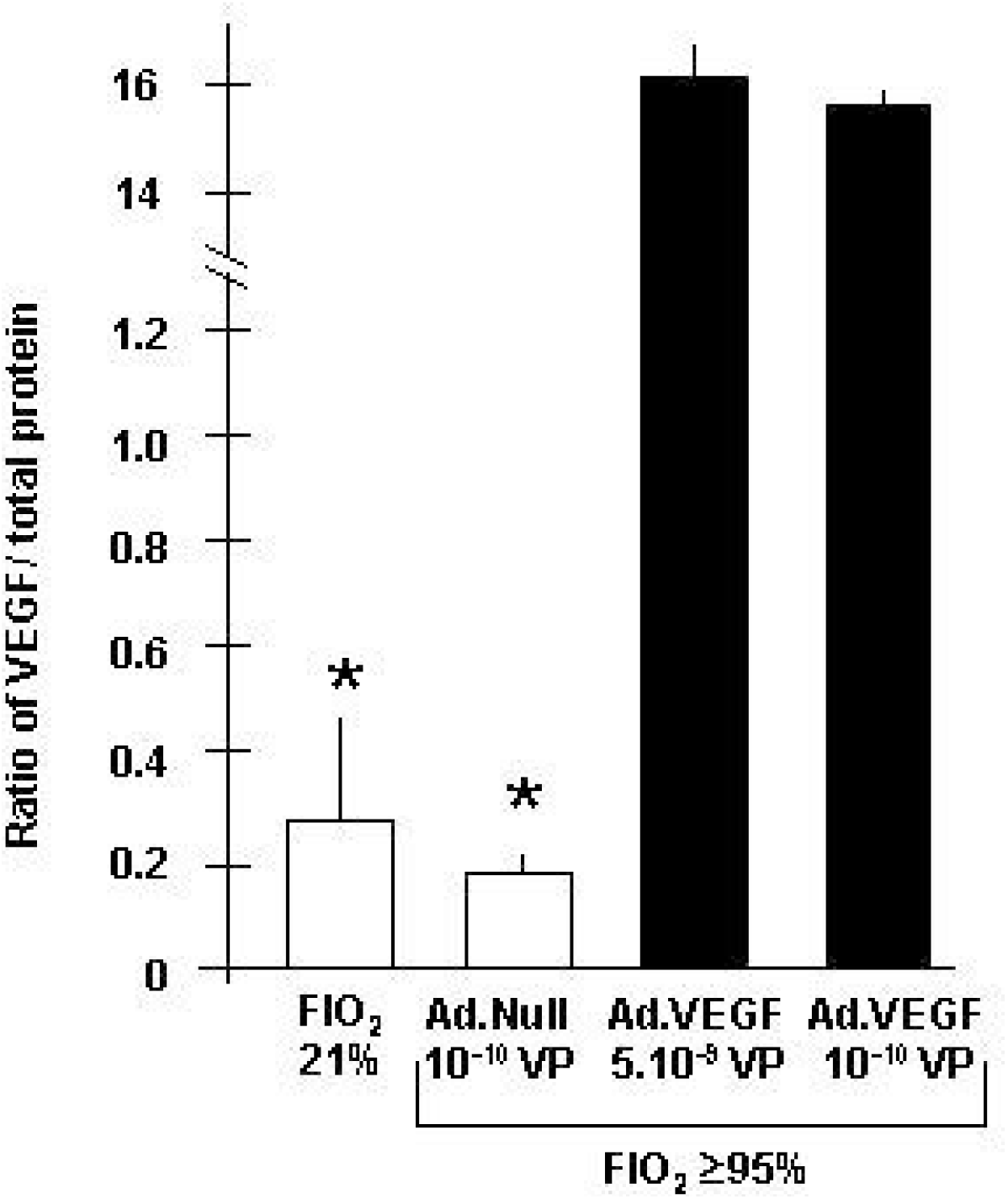
Efficiency of transduction. Concentrations of VEGF in lung homogenates from rats under room air (fraction of inspired oxygen [FIO_2_] 21%, open bar) and 48 h after exposure to high levels of hyperoxia (FIO_2_ ≥95%, solid bars). Adenoviral transduction was performed 48 h before starting oxygen exposure. The levels of VEGF were assessed using a commercial ELISA (n = 5 for each group). *: p<0.001 as compared to Ad.VEGF groups.

#### Assessment of lung injury

##### Lung histology

Hyperoxia induced classical airspace and interstitial injuries in rats pretreated with Ad.Null. After administration of 5.10^9^ or 10^10^ VP of Ad.VEGF, histological analysis depicted areas of patchy cell proliferation under normoxia, the sole difference between the two doses being an increased and diffuse recruitment of inflammatory cell with the higher dose of adenovirus. Under hyperoxic condition, a similar pattern of patchy areas of cell proliferation was observed together with inflammatory cell recruitment (figure 3), which was more marked with the higher dose of adenovirus. In normoxia, Ad.Null transduction was associated with no change in lung histology as compared with normal not-treated lung (data previously described by our laboratory [15]).

**Figure 3.**
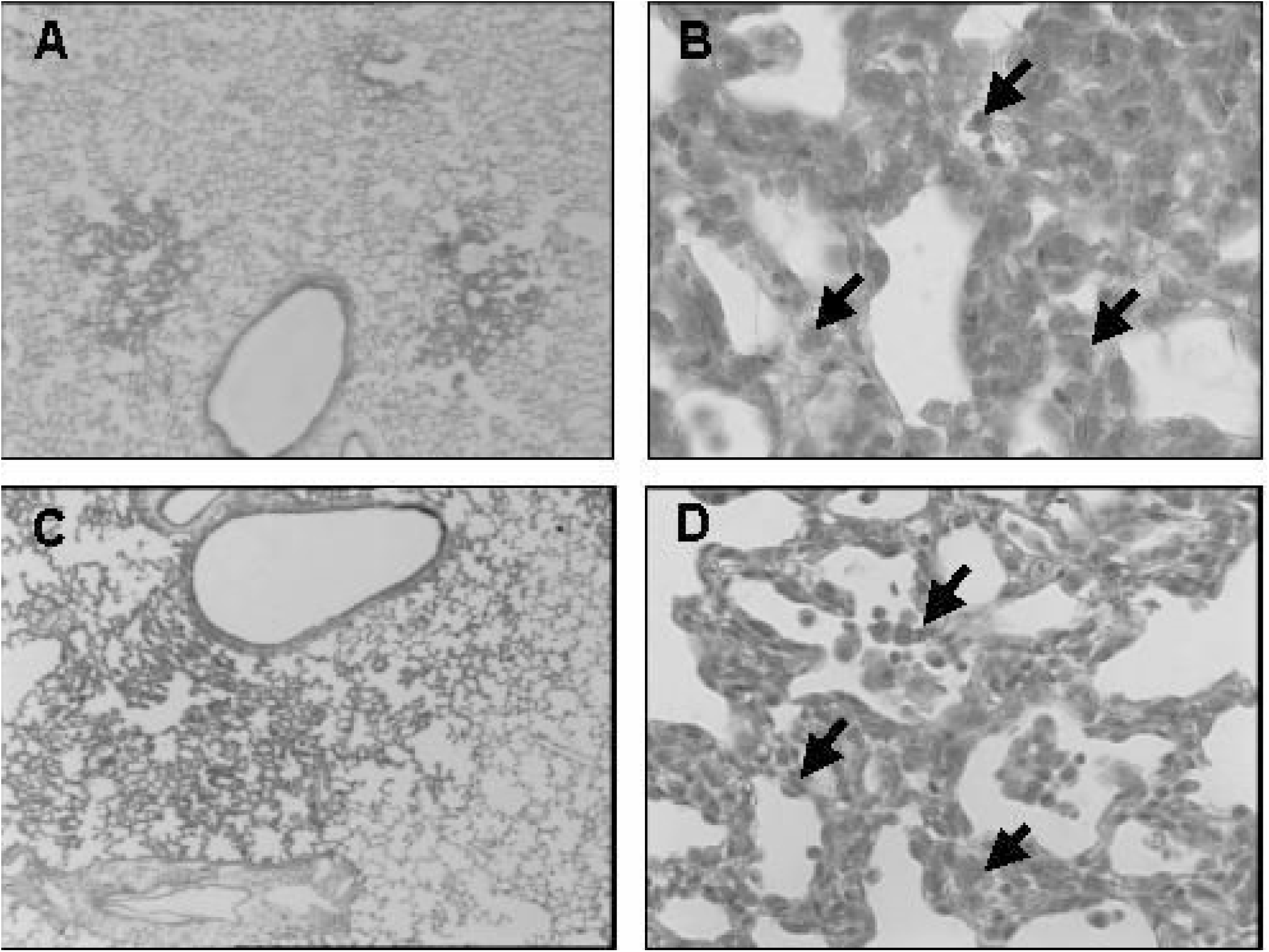
Histological lung sections of rats treated by Ad.VEGF (10^10^ VP). Four days after adenoviral transduction, abnormal hyperplastic areas are visible (panel A: normoxia, panel C: hyperoxia; original magnification x2.5); higher power views show hyperplasia with areas of edema and inflammatory cell infiltrate indicated by the black arrows (panel B: normoxia, panel D: hyperoxia; original magnification x25).

In lung sections, semi-quantitative evaluation of inflammation showed major recruitment of neutrophils and macrophages (alveolar and interstitial) in patchy areas of Ad.VEGF 10^10^ VP treated animals under both normoxia and hyperoxia. Ad.Null 10^10^ VP treated animals exhibited no inflammation under normoxia and moderate inflammatory cell recruitment (mainly neutrophils) under hyperoxia (figure 4). Overall, Ad.VEGF transduction depicted more inflammation under hyperoxia than control animals, which was mainly related to interstitial inflammation in localized areas of endothelial proliferation.

**Figure 4.**
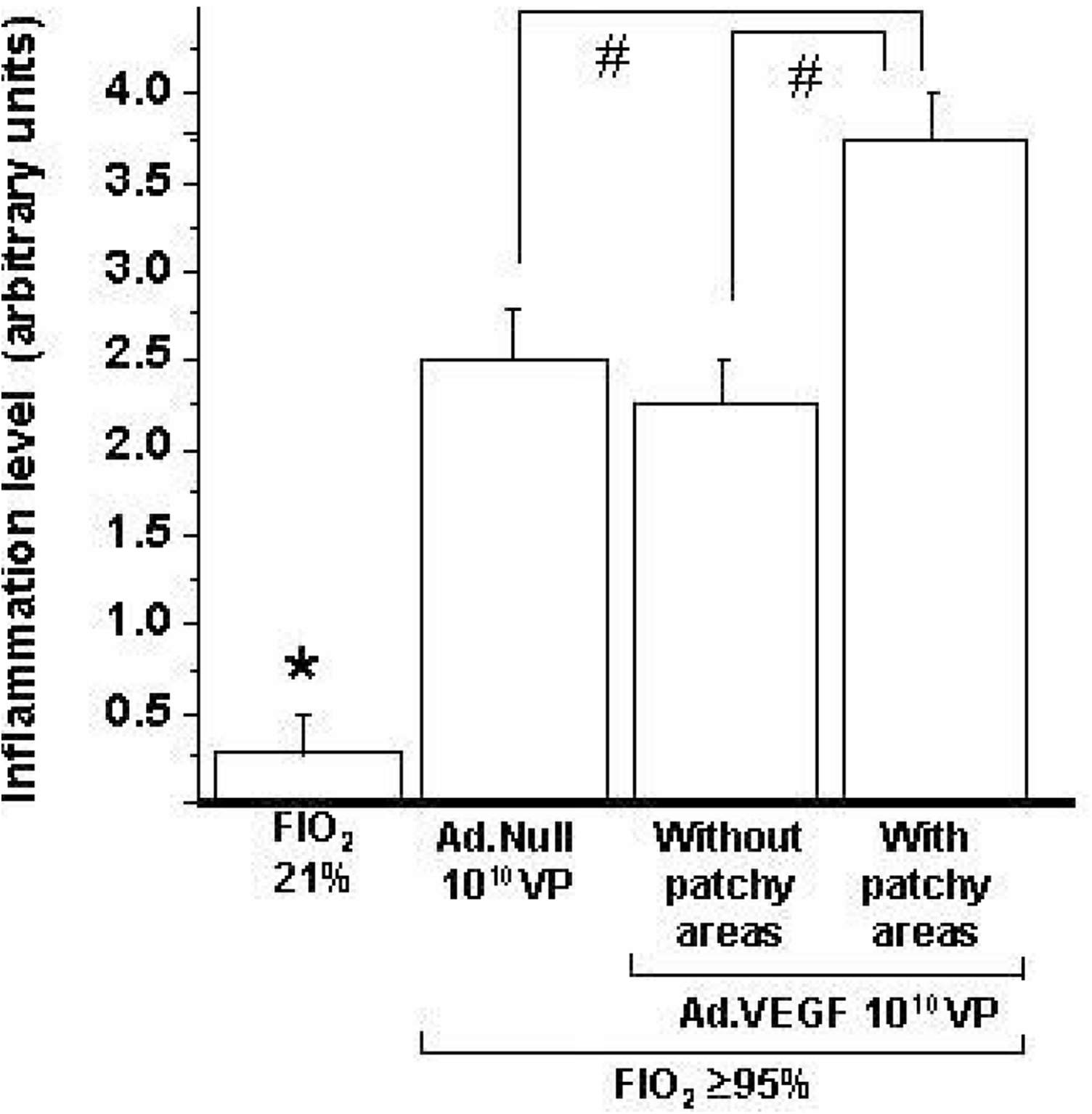
Quantification of inflammatory cell recruitment. Lung sections were examined in a blind manner to quantify inflammatory cell recruitment. Three tissue sections (10 fields in each section) were analyzed.

##### CD31 immunohistochemistry

Endothelial injury was suggested by the significant decrease in CD31 staining in lungs of rats from hyperoxic group challenged with Ad.Null as compared with sections from control rats under normoxia. The patchy areas of cell proliferation stained clearly for CD31. The quantification of the staining indicated that the decrease related to hyperoxia exposure was prevented by Ad.VEGF transduction even when patchy areas of endothelial proliferation were not taken into account (figure 5).

**Figure 5.**
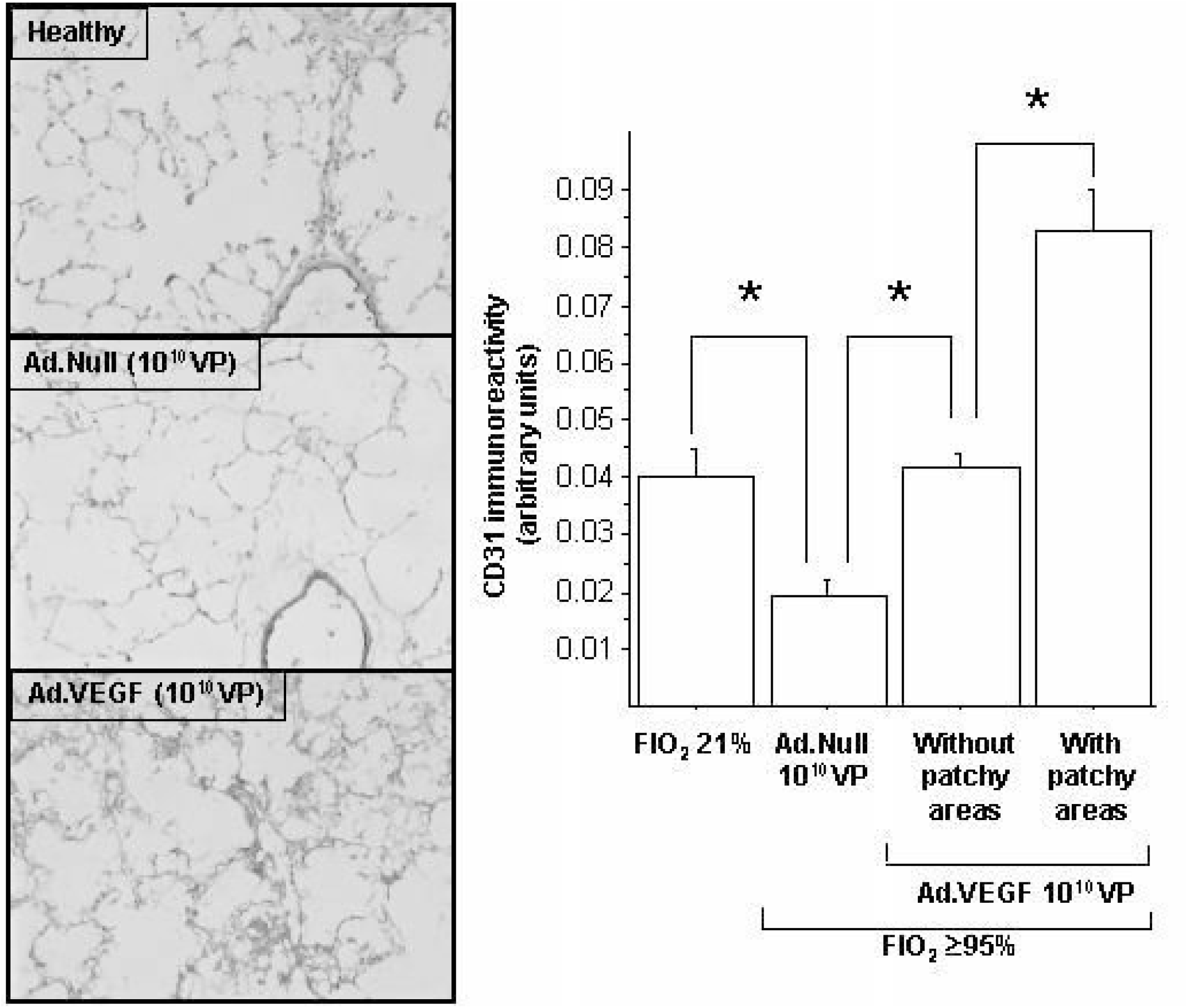
Quantification of microvascular density. Left panel: CD31 immunoreactivity in rat lungs. A decrease in CD31 staining was evidenced in Ad.Null rats under hyperoxia as compared to healthy animals whereas patchy areas of cell proliferation under VEGF transduction were clearly stained (original magnification x10). Control with primary antibody omitted showed no staining (data not shown). Right panel: Quantification of vascular density assessed by the ratio of CD31 immunostaining density/total tissue area in lung sections with Perfect Image software (Clara Vision, Orsay, France). Tissue sections (n=3 per animal) analysis of 10 fields of 4 animals in each group. * P <0.05.

##### Quantification of apoptotic cells

Apoptosis staining was very low in normoxic non-treated rats. Hyperoxic rats treated with Ad.Null and hyperoxic rats treated with Ad.VEGF (whatever the type of quantified area) exhibited significant difference as compared with control group in normoxia (figure 6). Apoptotic cells were both localized into alveoli (inflammatory cells) and alveolar capillary wall (epithelial and endothelial cells). Apoptosis staining in patchy areas was significantly lower than staining in non-patchy areas, suggesting a local inhibitory role of VEGF on endothelial apoptosis.

**Figure 6.**
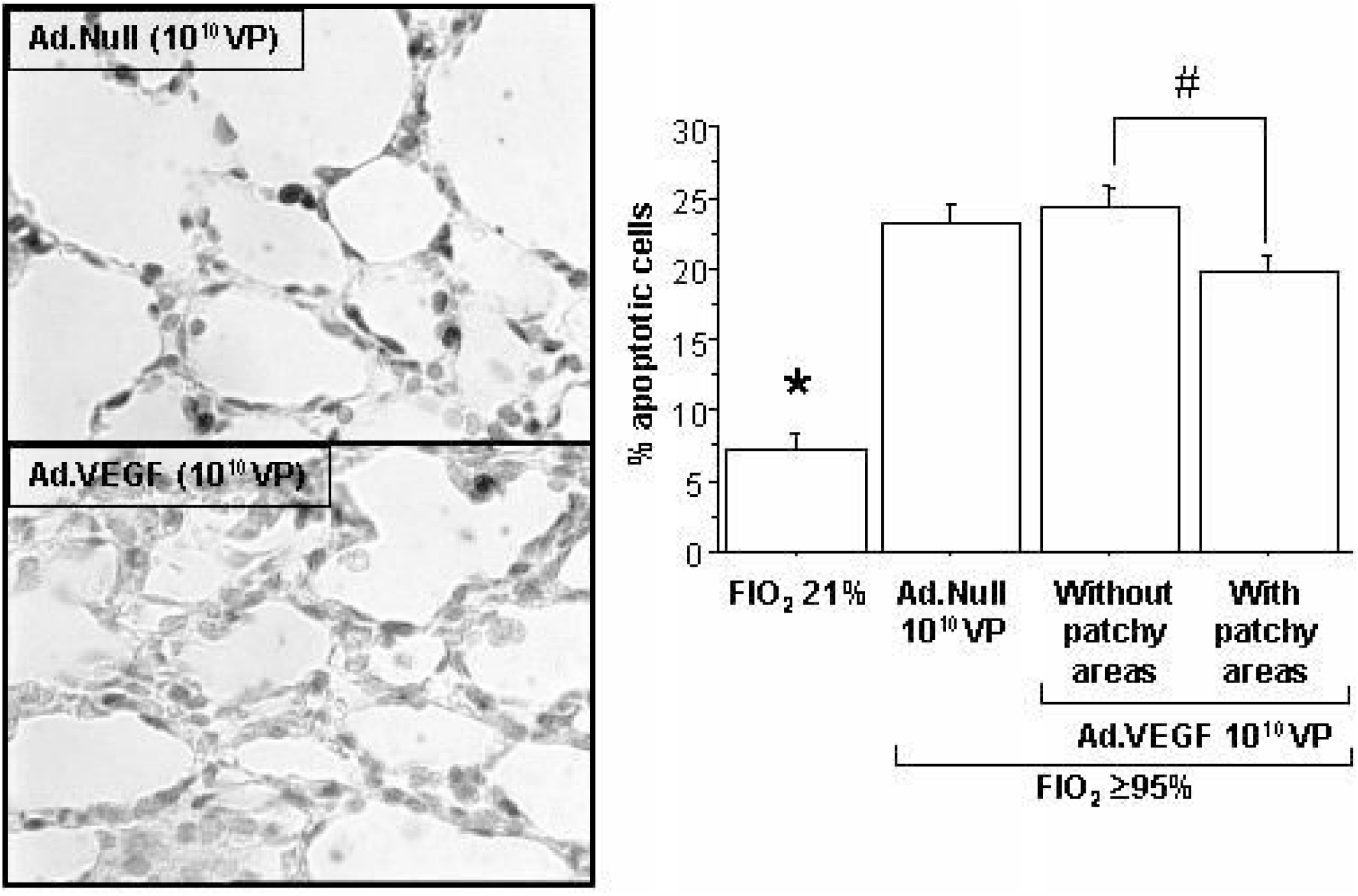
Quantification of apoptotic cells. Left panel: TUNEL staining in rat lungs. Rats were instilled with Ad.Null 10^10^ VP or Ad.VEGF 10^10^ VP and subsequently (2 days later) exposed to hyperoxia (original magnification x40). Black arrows indicate dark brown nuclei of apoptotic cells. Control with TdT reaction omitted showed no staining (data not shown). Right panel: Quantification of apoptosis (% of apoptotic cells / total cell). Analysis of 10 fields per tissue section was performed (n=4 rats for each group). * P <0.0001 as compared to other groups; #: P <0.05.

##### Quantification of alveolar inflammation

To further differentiate the respective roles of VEGF transduction and hyperoxia-injury, alveolar cell recruitment was analyzed in rats under normoxia and hyperoxia (figure 7). A similar pattern of alveolar inflammation was observed under hyperoxia in animals treated with either Ad.Null or Ad.VEGF. Under normoxia, a dose effect relationship on alveolar inflammation was observed with Ad.VEGF.

**Figure 7.**
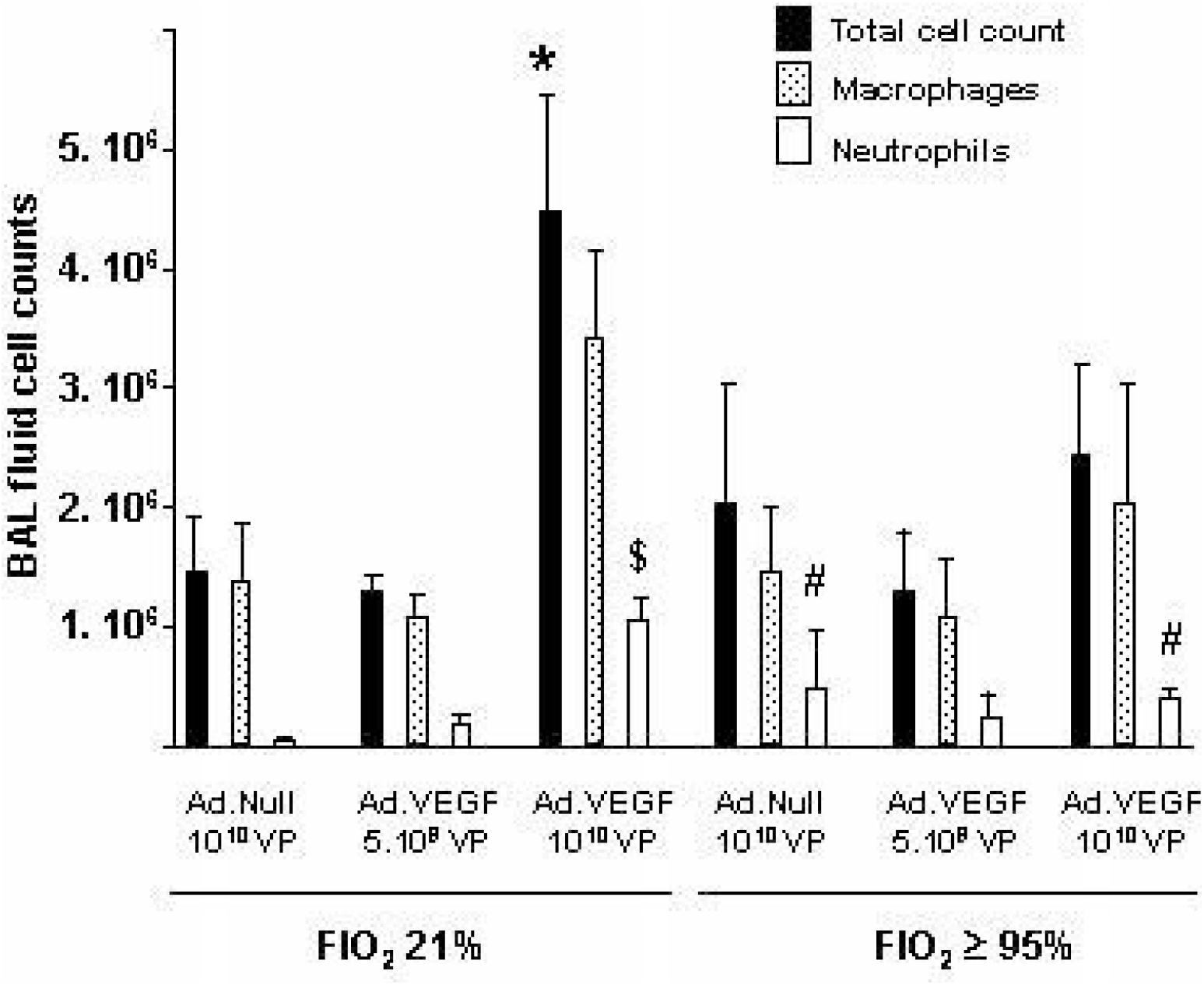
Quantification of alveolar inflammation. Inflammatory cells in bronchoalveolar lavage [BAL] fluid of rats 4 days after intratracheal Ad.VEGF (5. 10^9^ VP or 10^10^ VP) or Ad.Null (10^10^ VP) administration under normoxia and hyperoxia. Inflammatory cells in BAL fluids were recovered by cytospin centrifugation and identified by differential staining (n = 5 for each group). * P<0.005 as compared to Ad.Null values, ^§^ P<0.05 as compared to Ad.Null values, ^#^ P<0.05 as compared to Ad.Null values in normoxia.

##### Wet-to-dry lung weight ratio

Under normoxia, transduction with Ad.VEGF induced lung edema as compared to Ad.Null transduction. Under hyperoxia, lung edema was observed in control rats (Ad.Null) whereas it did not further increase in rats challenged with Ad.VEGF (figure 8).

**Figure 8.**
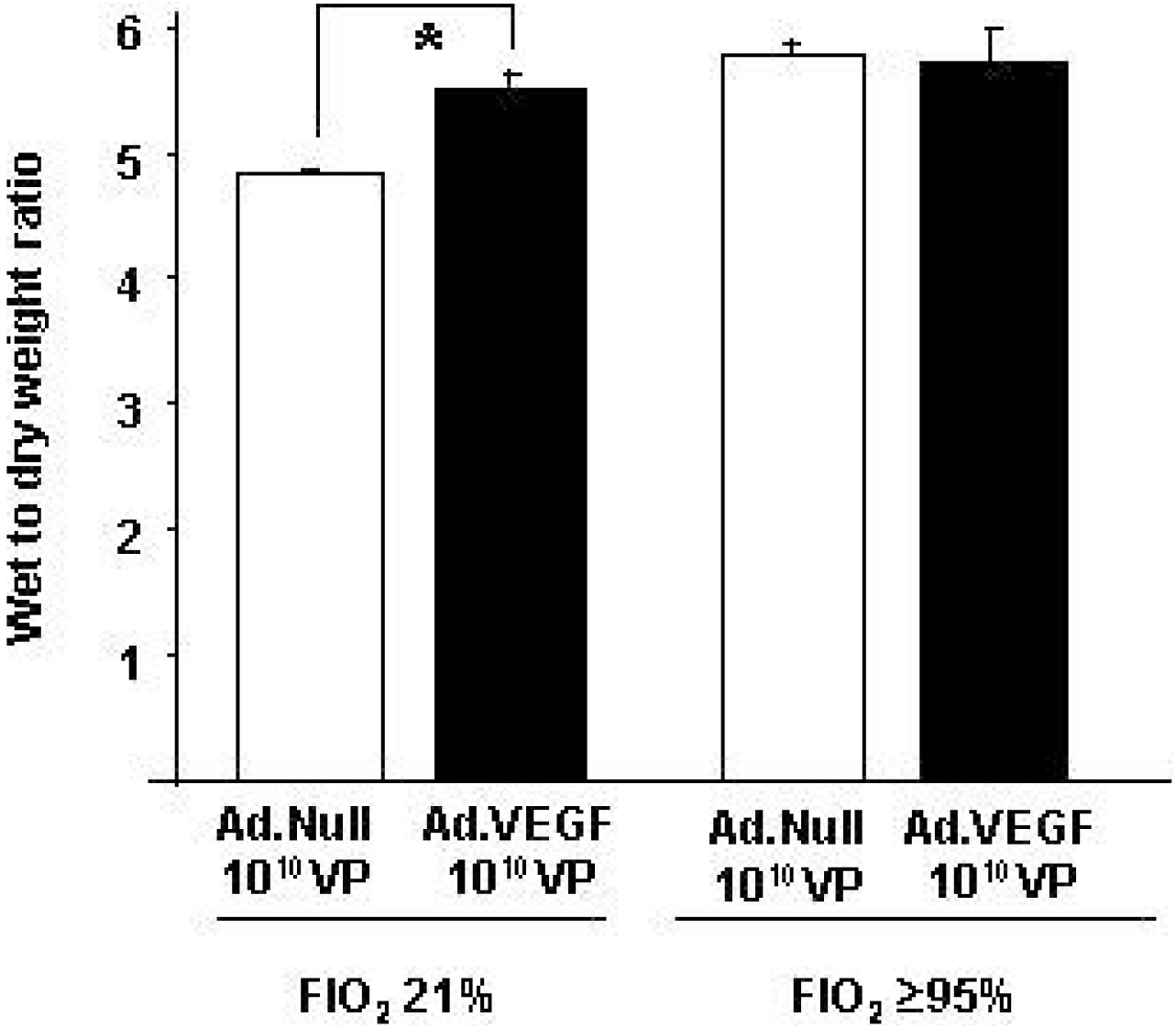
Quantification of lung edema. Effect of administration of Ad.VEGF 10^10^ VP or Ad.Null 10^10^ VP (control vector) on wet to dry weight ratio under 21% or ≥95% oxygen. Lungs were harvested two days after air or oxygen exposure and proceed as described in Methods (n = 5 for each group). * P d 0.05.

## Discussion

The role of VEGF and related molecules in acute lung injury/ARDS remains controversial as stated in a recent review [1]. In this experimental study, we tried to explore its role in hyperoxia-induced lung injury. Transduction of VEGF-A was effective since a dramatic increase in concentration of the growth factor was evidenced in lung homogenates, an inhibition of endothelial loss was observed in lung tissue and since obvious endothelial proliferation was evidenced in some areas. These results are consistent with those of Factor and colleagues who demonstrated that acute hyperoxic lung injury does not impede adenoviral-mediated alveolar gene transfer [16]. Ad.Null was considered in our study as a reference similar to non-treated group on the basis of our previous works since adenovirus vector without transgene used is not deleterious [11, 15]. Hyperoxia-mediated lung injury results in both endothelial and epithelial injuries, the former being predominant [10]. Similarly to epithelial injury, endothelial injury during hyperoxia may be related to both apoptosis and necrosis [17]. It is difficult to evaluate their respective contributions in vivo. In our model we found a diffuse increase in apoptotic cells (TUNEL) under hyperoxia as compared to normoxia, which is in agreement with the recent study of He and colleagues [18]. The increase in apoptosis was milder in the areas of endothelial proliferation that is probably related to a local effect of VEGF transduction. It has to be noted that only one marker of apoptosis was used, which constitutes a limitation of this study. Hyperoxia-induced injury led to a decrease in vascular bed as previously evidenced [10]. VEGF transduction prevented this decrease even when areas of patchy cell proliferation were not taken into account for the quantification, arguing for a role of apoptosis-induced diffuse vascular injury in this model. Along this line, we previously demonstrated a negative relationship between decreased VEGF concentration and endothelial apoptosis in lung tissue of patients with ARDS [8]. The patchy areas of endothelial proliferation together with local inhibition of apoptosis were probably related to supra physiologic local concentrations of VEGF due to focal transgenic expression.

Despite VEGF overexpression and its functional effects in this model, only a mild effect was evidenced on survival. Along this line, it has to be noted that the effect of VEGF inhibition during IL-13 overexpression in the study of Corne and colleagues had only a marginal effect on survival [19], and a moderate increase in survival under VEGF overexpression has been found in the recent study of He and colleagues [18]. Overall, the role of endothelial survival as a critical factor for animal survival is still debated [20].

The two doses of VEGF transduction that were used led to a similar increase in VEGF concentration that did not lead to a similar increase in survival. This may mimic the in vitro condition where expression increases in a manner dependent on the dose of adenovirus administered up to a plateau. At doses of adenovirus above this plateau, expression does not increase further and there is evidence of cytotoxicity related to the virus. Increasing the dose of adenovirus from 5.10^9^ to 1.10^10^ resulted in no further increase in VEGF expression but did result in toxicity as evidenced by alveolar neutrophilia. In aggregate, the sole difference that was observed between the two doses of VEGF transduction was the degree of alveolar inflammation (diffuse inflammation with the high dose versus localized to areas of endothelial proliferation with the low dose). This difference was only clearly evidenced under normoxia, whereas it was not significant under hyperoxia that modified inflammatory response (inhibitory effect of VEGF induced-inflammation). This difference, associated with the inhibition of loss of endothelial cells, may explain the respective effects of the two doses on survival. The effects of neutrophil influx during acute lung injury remains debated. On one hand, both Ad.heme oxygenase-1 administration and extracellular superoxide dismutase overexpression have been associated with reduction in neutrophil influx toward alveolar space and enhanced survival [21, 22]. On the other hand, several measures inducing lung inflammation such as various cytokine overexpression have previously been associated with an increase in survival during hyperoxia-induced lung injury [19, 20, 23]. Interestingly, two articles have recently demonstrated a pro-inflammatory effect of VEGF [24, 25].

In patients with ARDS we and others found a decrease in VEGF concentration in bronchoalveolar lavage fluid [6, 7], and more recently in lung tissue [8]. Hyperoxia-induced lung injury has consistently been associated with a decrease in lung VEGF mRNA [26-28]. During hyperoxia, VEGF protein can decrease in tissue while its concentration in epithelial lining fluid depends on the time of assessment [27, 28]. In our study, the decrease in VEGF protein concentration under hyperoxia in lung tissue was not significant, which may reflect this complex time-dependency. Our results do not suggest that VEGF will constitute a promising rescue therapy during ARDS.

In conclusion, we demonstrate that despite evidence of functional consequences such as a decrease in injury to vascular bed, lung VEGF-A transduction has a marginal effect of on animal survival during hyperoxia-induced injury.

## Acknowledgments

This work was supported by Leg Poix (Académie de Paris), by Collège des Professeurs de Pneumologie (France) and by Fondation Recherche Médicale (supported grants for W. Raoul).

## Authors’ contributions

WR and VL carried out the animal experiments and assays, and drafted the manuscript. SE and SA participated in the design of the study and performed the statistical analysis. CD2 and BM conceived of the study, and participated in its design and coordination and helped to draft the manuscript. All authors read and approved the final manuscript.

